# Using machine learning to detect coronaviruses potentially infectious to humans

**DOI:** 10.1101/2022.12.11.520008

**Authors:** Georgina Gonzalez-Isunza, M. Zaki Jawaid, Pengyu Liu, Daniel L. Cox, Mariel Vazquez, Javier Arsuaga

**Affiliations:** University of California, Department of Microbiology & Molecular Genetics, Davis, CA, USA; University of California, Department of Molecular & Cellular Biology, Davis, CA, USA; Department of Mathematics, University of California, Davis, CA, USA; Department of Physics, University of California, Davis, USA

## Abstract

Establishing the host range for novel viruses remains a challenge. Here, we address the challenge of identifying non-human animal coronaviruses that may infect humans by creating an artificial neural network model that learns from the binding of the spike protein of alpha and beta coronaviruses to their host receptor. The proposed method produces a human-Binding Potential (h-BiP) score that distinguishes, with high accuracy, the binding potential among human coronaviruses. Two viruses, previously unknown to bind human receptors, were identified: Bat coronavirus BtCoV/133/2005 (a MERS related virus) and *Rhinolophus affinis* coronavirus isolate LYRa3 a SARS related virus. We further analyze the binding properties of these viruses using molecular dynamics. To test whether this model can be used for surveillance of novel coronaviruses, we re-trained the model on a set that excludes SARS-COV-2 viral sequences. The results predict the binding of SARS-CoV-2 with a human receptor, indicating that machine learning methods are an excellent tool for the prediction of host expansion events.

## Introduction

Most novel viral human diseases, particularly those that have caused recent epidemics, are known to have originated in non-human animal hosts^1,2,3^. Host expansion, the ability of a virus to cross species, is an essential step in the evolution of such viruses^3,4,5^. COVID-19 is a recent example of a disease caused by a host expansion event that permitted SARS-CoV-2, a SARS-related coronavirus, to propagate from a yet unknown non-human animal to humans^5^. Alpha and beta coronaviruses affect a wide range of animals interacting with humans, including farm animals and camels, thus facilitating zoonotic transmission^6,7^. Moreover, all seven human coronaviruses belong to either the alpha or beta coronavirus genus^7^. While several studies have confirmed bats and rodents as natural hosts for the alpha and beta coronaviruses affecting humans, there is evidence of intermediate hosts that facilitate evolutionary events, leading to strains that eventually propagate in humans^1,6,8^. Determining which non-human animal viruses may infect humans remains a challenge.

Experimental evidence is still the gold standard used to determine whether a virus can infect a host^9,10^. However, the complete host range of a virus is often unknown. Recent studies have used diverse in-silico techniques to predict viral hosts and host expansion events, including qualitative expert analysis^11^, probabilistic^12^ and machine learning (ML)^13, 14, 15, 16, 17^ models.

The problem of host prediction is commonly addressed using similarity analysis of viral genomes, where similar genomes are more likely to share the same hosts^10,18^. Host prediction through genome similarity can be achieved by alignment-based or alignment-free approaches^17,19^. Computational efficiency of alignment-based approaches decreases with the product of the lengths of the sequences being aligned^19,20^ and are sensitive to genome rearrangements^19,20,21^. These observations suggest alignment-free approaches may be preferred when datasets are very large or sequences in the dataset are the product of recombination events. However, most alignment-free approaches disregard the relative position of the residues along the sequence^14^.

Some alignment-free studies aimed at predicting the host of a specific species of virus^13,14^, while others^15,16,17^ created models to uncover signals common to different viruses (*e.g*. Zika, influenza, coronavirus) affecting a large group of hosts such as Chordata (vertebrates and others)^15,17^. Although common signals between completely different families of viruses are useful for host prediction, these studies include only a limited number of representatives of each taxa across hosts and disregard the specific properties of the virus, preventing further mechanistic analysis of host expansion pathways.

In this work, we study the potential of alpha and beta coronaviruses to cause human infection. In particular, we aim at predicting whether the spike (S) protein of a coronavirus binds a human receptor. The S protein decorates the exterior of the viral envelope and is key in host expansion since its binding to the host receptor protein triggers the infection process^22,23,24^. Starting with a collection of amino acid sequences from the S protein, we build a machine learning model that predicts binding to a human host receptor. We propose a skip-gram model which uses a neural network to transform the data into vectors. These vectors encode the relationship between neighboring protein sequences of length k (*i.e*. k-mers). A classifier uses these vectors to score each sequence according to its binding potential to a human receptor. We call this score the human-binding potential (h-BiP). We use a dataset consisting of 2,534 unique spike sequences from alpha and beta coronaviruses spanning all clades and variants (see Methods). The classifier is highly accurate, and its h-BiP score is highly correlated with sequence identity against human viruses. Moreover, the proposed h-BiP score also discriminates the binding potential in cases with similar sequence identity and detects binding in cases of low sequence identity. We identify two viruses, Bt133^25^ and LYRa3^26^, with high h-BiP values and yet unknown human binding properties. Consistent with this finding, a phylogenetic analysis shows that Bt133 and LyRa3 are related to non-human viruses known to bind human receptors. Furthermore, a multiple sequence alignment of the receptor binding motifs (RBM) of Bt133 and of LYRa3 with their related viruses revealed that they conserve the contact residues with the human receptor. Molecular dynamics (MD) of the receptor binding domain (RBD) validates binding and identifies contact residues with human receptors. Finally, we test whether this model can be used for the surveillance of host expansion events. We emulate the conditions prior to SARS-CoV-2 emergence by excluding all SARS-CoV-2 sequences from the training set and find that the re-trained model predicts binding of the wild type of SARS-CoV-2 to a human receptor.

## Results

### h-BiP: a machine learning approach for scoring human-binding potential of coronavirus sequences

We propose a human-Binding Potential (h-BiP) score that assigns a value between 0 and 1 to spike proteins of alpha and beta coronaviruses. The outline of the method to obtain h-BiP is shown in Fig. 1, and a full description is available in Methods section. First, we partitioned each protein sequence as a sequence of trimers, which we used to produce trimer embeddings in a large-dimensional Euclidian space. We selected a skip-gram model^27^ to ensure that each trimer embedding was informed by neighboring trimers within a context window. Next, an embedding for the entire protein sequence was generated by adding all of its trimer embeddings. Each coronavirus was labeled as positive or negative according to its published binding annotation to a human receptor (Table 1). To produce the h-BiP score, we built a logistic regression classifier on the sequence embeddings to predict binding. Viruses with h-BiP score of 0.5 or higher were classified as likely to bind a human receptor.

**Fig. 1:**
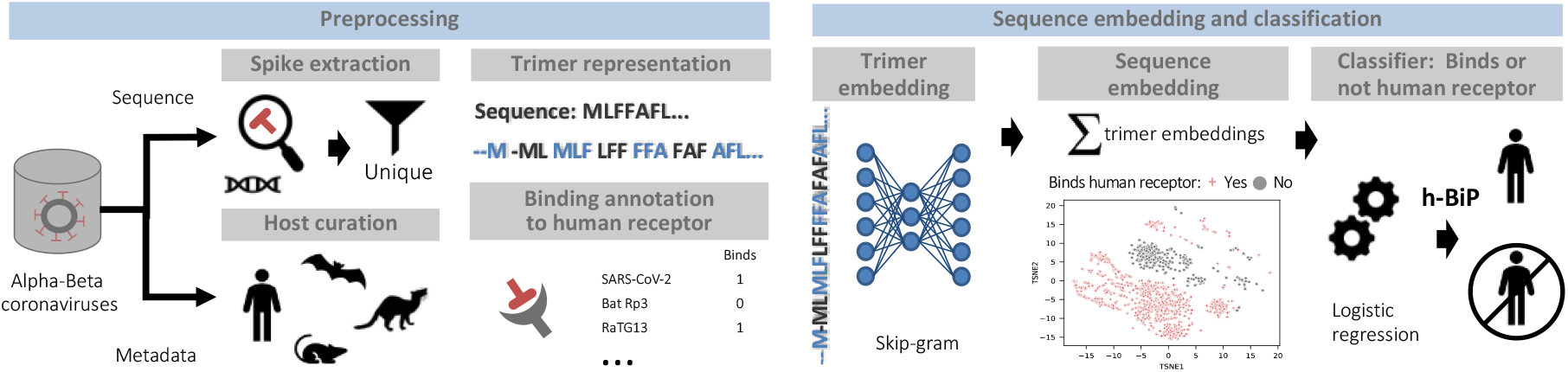
Methodological workflow of the human-Binding Potential (h-BiP) score. **LEFT:** Preprocessing sequences from alpha and beta coronaviruses. **Top:** Whether the S protein was available from annotation or by extraction from whole-genome, the dataset consists of 2,534 unique S protein sequences. Each protein sequence is transformed into a trimer (3 amino acid) representation by sliding a window one amino acid at a time. **Bottom:** We curated the host field and annotated the sequences according to their binding status to human receptors. Regardless of the host, a virus is considered positive for binding if there is experimental evidence of binding to a human receptor. **RIGHT:** A skip-gram model uses a neural network to generate trimer embeddings of a fixed dimension (d=100). These trimer embeddings are numerical vectors that encode information from all neighboring trimers within a context window in the protein sequence. Next we compute the final sequence embedding (d=100) by adding up all of its trimer embeddings. The scatterplot shows a visualization for the embeddings from all viruses after using t-distributed stochastic neighbor embedding (tsne) to reduce dimensionality. Finally, all sequence embeddings feed a classifier (logistic regression) to produce the h-BiP score that learns from the binding information of alpha and beta coronaviruses. An h-BiP score greater than or equal to 0.5 flags the virus as likely for human binding.

**Table 1:** Viruses from non-human hosts according to experimental evidence of binding.

| Binds to human receptor |
| --- |
| RaTG13<sup>31, 35</sup>, LYRa11<sup>34, 35</sup>, WIV1<sup>8, 35, 38</sup>, WIV16<sup>8, 38</sup>, RsSHC014<sup>34, 35</sup>, Rs4231<sup>34</sup>, Rs4084<sup>34, 39</sup>, Rs4874<sup>40</sup>, Rs7327<sup>40</sup>, Rs4231<sup>40</sup>, Khosta-1<sup>41</sup>, Khosta-2<sup>41</sup>, Ty-HKU4<sup>24, 33</sup>, PCoV_GX<sup>31</sup>, PCoV_GD<sup>31</sup>, A022G<sup>38</sup>, SZ03<sup>38</sup>, B039G<sup>38</sup>, DcCoV UAE-HKU23<sup>42</sup>, Bovine coronavirus isolate Alpaca<sup>32</sup>, camel alphacoronavirus / hCoV229E<sup>29</sup>, MERS Middle East strain<sup>28</sup>, HKU25<sup>33</sup>, YN2018B<sup>39</sup>, Rs9401<sup>39</sup>, Rs3367<sup>39</sup> |
| No evidence of binding to human receptor |
| Rc-o319<sup>35</sup>, Bat CoVZXC21<sup>35</sup>, ZC45<sup>34</sup>, HKU5<sup>24, 33</sup>, Bat Rf1<sup>35</sup>, Bat Rf4092<sup>35</sup>, Bat BM48-31<sup>34, 35</sup>, Bat Rp3<sup>35, 44</sup>, bat SL CoV As6526<sup>44</sup>, Bat Rm1<sup>40</sup>, As6526<sup>34</sup>, Yunnan 2011<sup>34</sup>, Shaanxi 2011<sup>34</sup>, Bat 279-2005<sup>34</sup>, Rs4237<sup>34</sup>, Rs4081<sup>34</sup>, Rs4247<sup>34</sup>, Bat HKU3-1<sup>35</sup>, HKU3-8<sup>34</sup>, HKU3-13<sup>34</sup>, GX2013<sup>34</sup>, Longquan-140<sup>34</sup>, YN2013<sup>34</sup>, Rf4092<sup>34</sup>, JL2012<sup>34</sup>, HuB2013<sup>34</sup>, HeB2013<sup>34</sup>, 273-2005<sup>34</sup> |

### The h-BiP score is highly accurate in predicting binding to human receptors for alpha and beta coronaviruses

We split the data into training (85%) and testing sets (15%) stratified by the following groups: hCoV-OC43, hCoV-HKU1, MERS, SARS-CoV-1, SARS-CoV-2, hCoV-NL63, hCoV-229E, other MERS-related, other Sarbecovirus, other Betacoronavirus, porcine epidemic diarrhea virus, other Alphacoronavirus.

Table 2 shows the prediction results for the full data set. The method achieved 99.5% accuracy, 99.6% sensitivity and 98.4% specificity in the test data set (see Supplementary Fig. 1). A total of 820 sequences had an h-BiP score less than 0.5. Of these, 813 were true negatives. While binding status was unknown for most of these sequences (766 out of 820), all 47 sequences (including variants) for which experimental studies found no evidence of binding to a human receptor were confirmed as non-binding by h-BiP (i.e. h-BiP score < 0.5). Sixteen viruses with unknown binding status had an h-BiP score ≥ 0.5, suggesting they may potentially bind human receptors. Fourteen of these sixteen viruses were MERS-related viruses from African dromedary camels. All MERS-related viruses of dromedary camels in the dataset (n=80) had an h-BiP score ≥ 0.5. Human transmission of MERS-CoV strains in dromedary camels from the Arabian Peninsula has been experimentally confirmed^28^, and some studies suggest that MERS-CoV strains from African camels may infect humans^29^. Two more viruses with unknown human binding status and a high h-BiP score were detected: Bat coronavirus BtCoV/133/2005 (Bt133) and Rhinolophus affinis coronavirus isolate LYRa3. Seven viruses were classified as false negatives (see Table 3). The h-BiP scores of these viruses ranged from 0.36 to 0.48, except Khosta-1 with a h-BiP=0.23.

**Table 2:** Confusion matrix for h-BiP on alpha and beta coronaviruses.

| h-BiP | Binds to human receptor |  | Total |
| --- | --- | --- | --- |
|  | Negative/Unknown | Positive |  |
| Negative<br>(h-BiP<0.5) | TN = 813<br>(47 neg, 766 unknown) | FN = 7 | 820 |
| Positive<br>(h-BiP≥0.5) | FP = 16<br>(unknown) | TP = 1698 | 1714 |
| **Total** | **829** | **1705** | **2534** |

True negatives (TN), false negatives (FN), false positives (FP) and true positives (TP)

**Table 3:** Viruses known to bind human receptors for which h-BiP is smaller than 0.5 (false negatives)

| Accession | Virus | Human receptor | h-BiP score |
| --- | --- | --- | --- |
| MZ190137 | Bat SARS-like coronavirus Khosta-1<sup>45</sup> | ACE2 | 0.226 |
| MZ190138 | Bat SARS-like coronavirus Khosta-2<sup>45</sup> | ACE2 | 0.360 |
| AWH65877 | Tylonycteris bat coronavirus HKU4 isolate CZ01 | dPP4 | 0.391 |
| ACJ35486 | Human enteric coronavirus 4408 | 9-O-acetylated sialic acid | 0.340 |
| ACT11030 | Human enteric coronavirus 4408 | 9-O-acetylated sialic acid | 0.346 |
| ABI93999 | Bovine coronavirus isolate Alpaca<sup>32</sup> | N-acetyl-9-O-acetylneuraminic acid | 0.396 |
| APD51499 | 229E-related bat coronavirus strain BtKY229E-8<sup>46</sup> | APN | 0.483 |

### The h-BiP score is consistent with sequence identity and expands viral classification

We verified that the proposed model, while consistent with sequence alignment results, provides additional information. Percent sequence identity (% identity) of a newly detected virus with known human viruses^10,18^ is often used to assess human infectivity. We computed the pairwise % identity between each of the 7 human coronaviruses and the S protein sequences in our dataset and selected the maximum for each sequence. All cases with 97 or higher protein % identity with known human coronaviruses had a h-BiP score greater than 0.5. Furthermore, the Pearson correlation between these maximum % identities and our proposed h-BiP score was 0.96 (Fig. 2). Hence, we conclude that our method is consistent with standard sequence identity approaches.

**Fig. 2:**
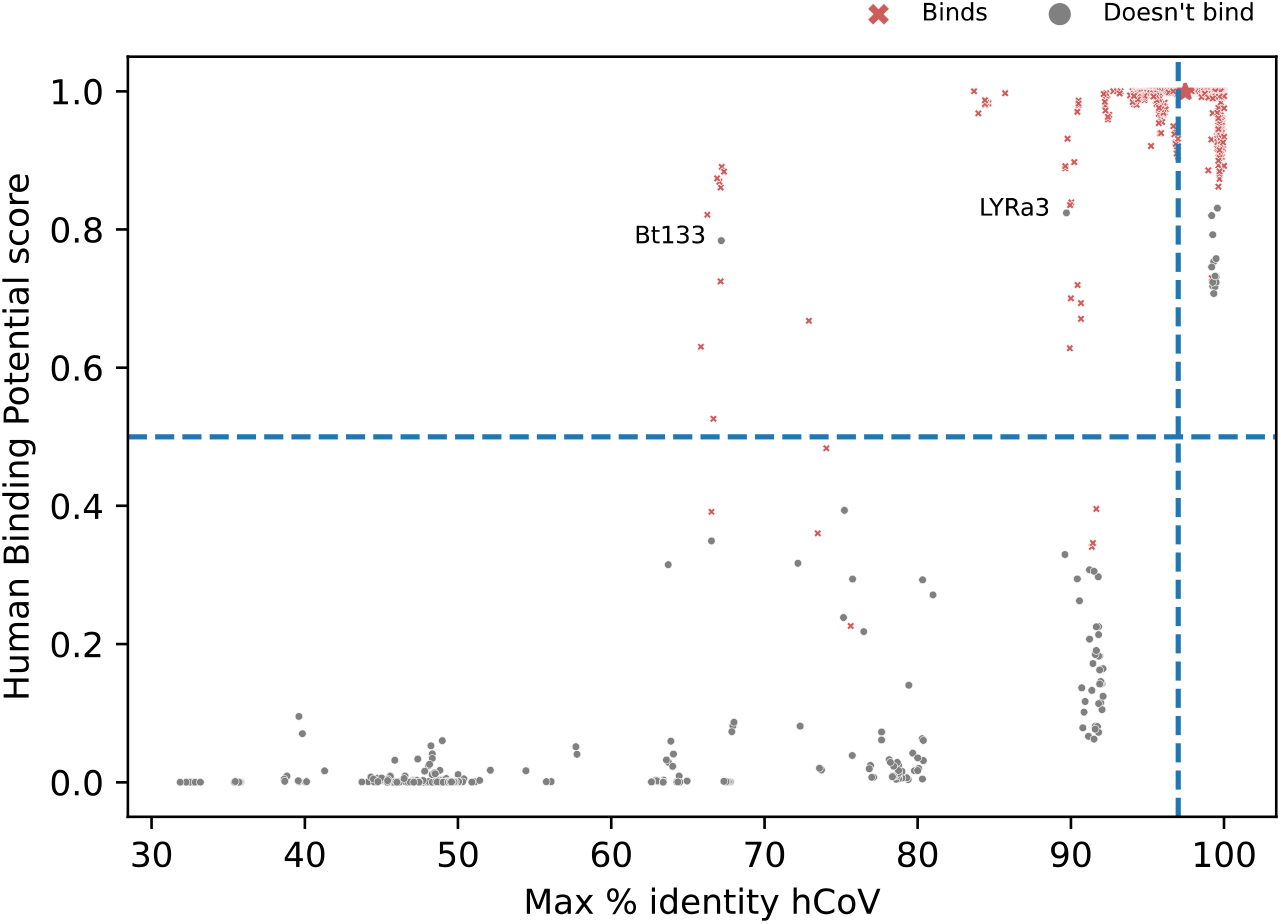
Comparison of sequence % identity and h-BiP score for alpha and beta coronaviruses. The x-axis represents the maximum % identity computed from a particular virus against the seven known human coronaviruses. The y-axis shows the h-BiP score. Each point in the graph represents a sequence in the dataset. Regardless of their host, red crosses depict sequences of viruses known to bind a human receptor, and grey points represent those viruses not known to do so (the latter including viruses for which in vitro studies found no evidence of binding). Points above the blue dashed horizontal line have a h-BiP score greater or equal than 0.5 (i.e. positive for binding). The blue dashed vertical line is the 97% identity reference line. The red rectangle contains the sequences of dromedary camels from African strains for which no human MERS cases have been reported. However, in vitro studies have shown binding to human receptors^29^. The spike protein of bat coronavirus RaTG13 (depicted with a red star) is known to bind to human receptor hACE2 and it has a 97.46% amino acid identity against SARS-CoV-2 and a 0.999 h-BiP score. Two viruses with h-BiP ≥ 0.5 and yet unknown binding: Bt133 and LYRa3 are highlighted.

Our study identified 12 bat viruses with maximum % identity <83% and an h-BiP ≥ 0.5. There is experimental evidence that these viruses bind to a human receptor, except for Bt133 (Table 1). A protein BLAST^30^ against all coronaviruses confirmed that Bt133 has a low % identity with any human coronavirus. The h-BiP score also discriminates between viruses that have similar % identity. For instance, the spike proteins from bovine and pangolin coronaviruses have % identities that range from 90% to 92% (See Fig. 2). The h-BiP scores for pangolin coronaviruses and bovine coronaviruses are greater than 0.97 and less than 0.4, respectively. This finding agrees with current experimental results that show binding of pangolin coronaviruses to human receptors^31^. To our knowledge, only the alpaca isolate from Bovine coronaviruses^32^ has been reported for human infections.

### The h-BiP score predicts that the S protein of viruses Bt133 and LYRa3 bind to human receptors

Our method assigned a h-BiP score>0.5 to Bt133 and LYRa3, suggesting that these two viruses with unknown human binding status may bind to a human receptor. Bt133 is a beta coronavirus from the Merbecovirus subgenus, and it is phylogenetically related to Ty-HKU4 (see Fig. 3a), a bat coronavirus for which there is experimental evidence of binding to human receptor dipeptidyl peptidase 4 (hDPP4)^24,33^. LYRa3 is a beta coronavirus from the Sarbecovirus subgenus, and it is phylogenetically related to *Rhinolophus affinis* coronavirus isolate LYRa11^26^ (see Fig. 3b), which binds human receptor angiotensin-converting enzyme 2 (hACE2)^34,35^. Host recognition and cell entry is mediated by the S protein^22,23,24^. The high sequence identity (97%) that the S protein of Bt133 shares with that of Ty-HKU4 suggests that Bt133 binds to hDPP4. Similarly, a 99% S protein sequence identity between LYRa3 and LYRa11 suggests that LYRa3 binds to hACE2.

**Fig. 3:**
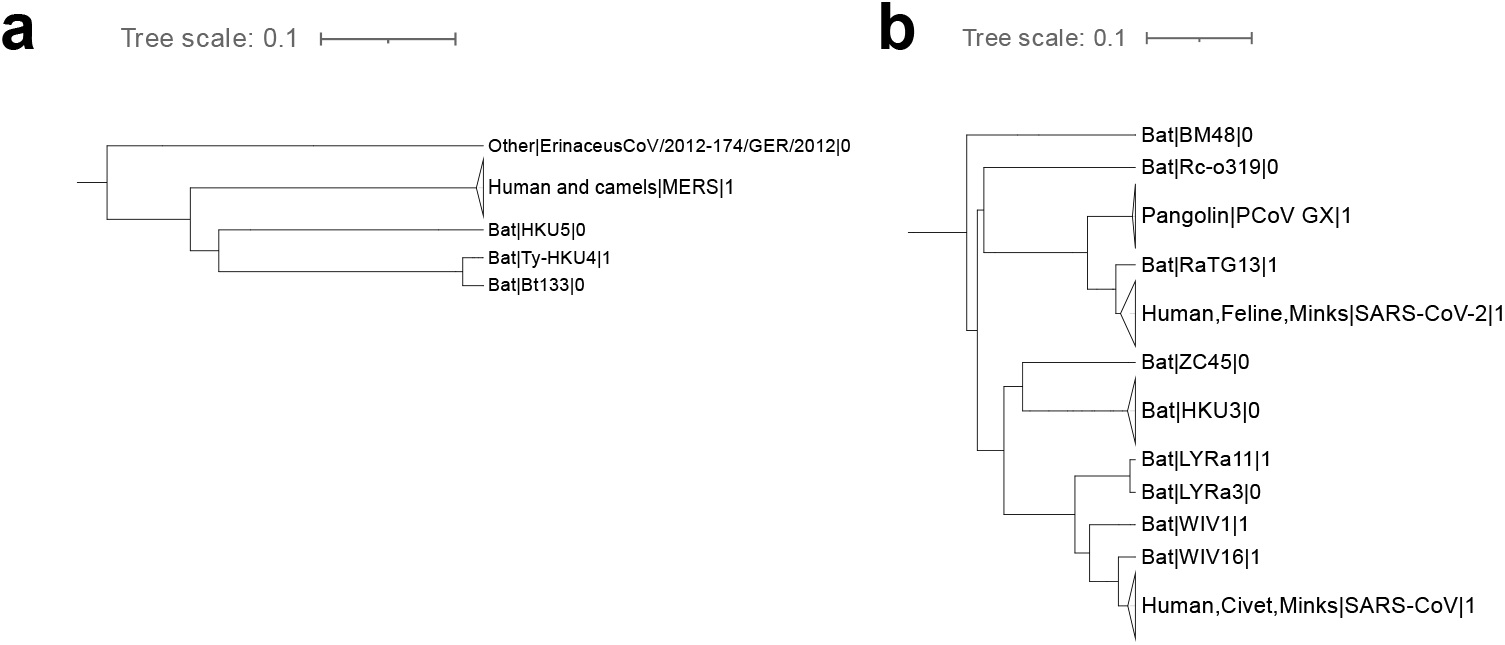
Phylogenetic tree for viruses related to Bt133 and LYRa3 at the S gene. Pruned version of maximum-clade-credibility tree generated from 424 alpha and beta coronaviruses (full tree available in Su pplementary Fig. 2). Each leaf shows the host, the name of the virus and the binding status separated by a pipe symbol. Non-human viruses with published binding annotation to a human receptor and human viruses have a binding status of 1 (0 otherwise). Solid gray triangles at the left of a leaf represent multiple variants in the particular leaf **a.** Phylogenetic tree for the Merbecovirus subgenus. Bat coronavirus Bt133 is phylogenetically related to Ty-HKU4. **b.** Phylogenetic tree for the Sarbecovirus subgenus. LYRa3 is phylogenetically related to LYRa11.

The S protein binds to the human receptor through the receptor binding domain (RBD). The RBD is composed of a core domain and the receptor binding motif (RBM) that comes in direct contact with the host receptor^24^. A multiple sequence alignment of Bt133 with typical members of the Merbecovirus subgenus at the RBM (See Fig. 4a) revealed that Bt133 conserves all 8 contact residues used by Ty-HKU4 to bind hDPP4^24,33^. In contrast, experimental studies from bat coronavirus HKU5^36^ did not find evidence of binding to hDPP4^24,33^. When compared to Ty-HKU4, HKU5 shows two deletions at the RBM and different amino acids at the contact residues of Ty-HKU4. Thus suggesting that Bt133 binds hDPP4.

**Fig. 4:**
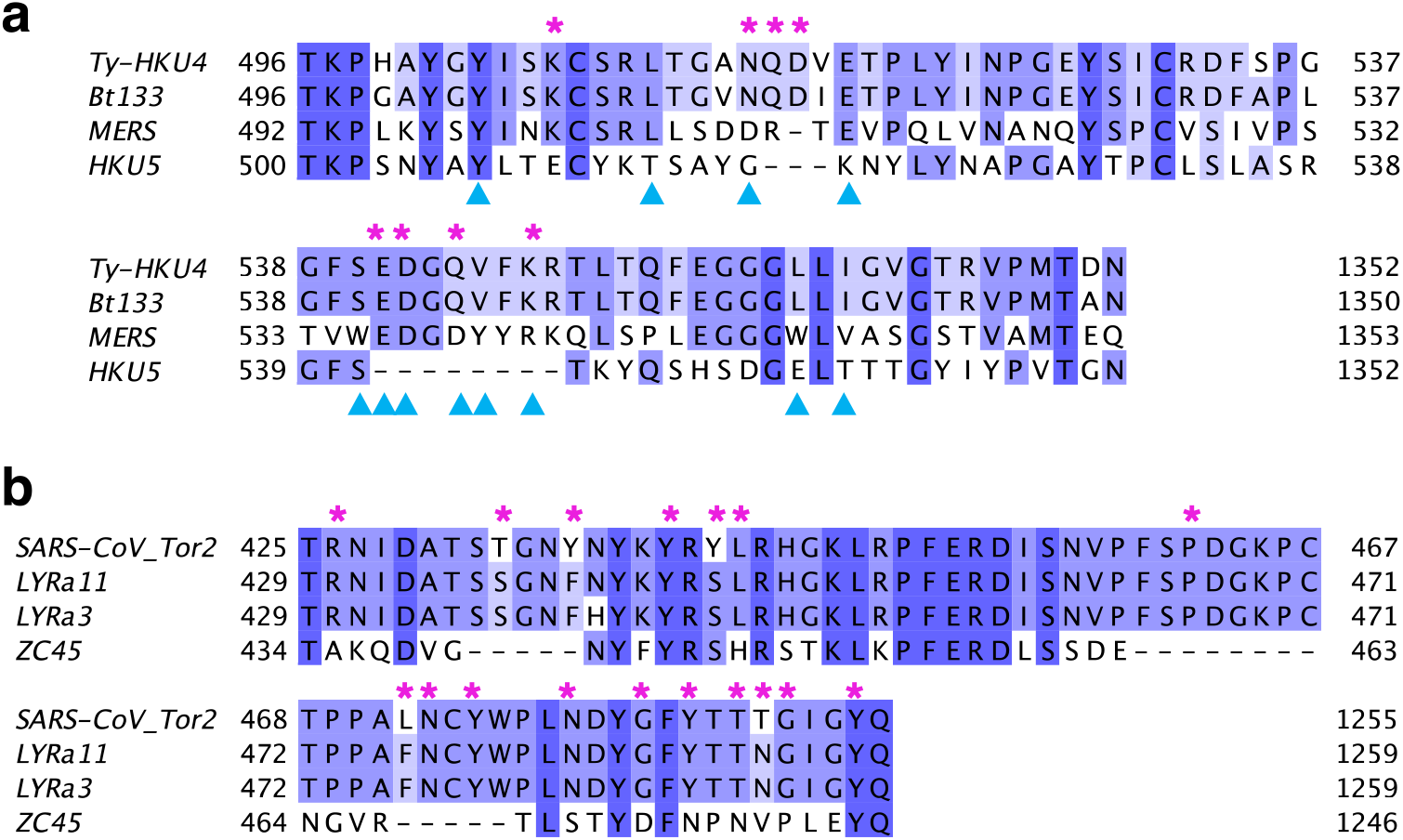
Multiple sequence alignment for phylogenetically related viruses at the RBM. Multiple sequence alignment was performed with MUSCLE^47^ and produce visualizations with Jalview^48^. A darker shade shows residues conserved in at least 50% of the sequences. **a**. Comparison of viruses related to Bt133 within the Merbecovirus subgenus. Ty-HKU4 and MERS are viruses known to bind human receptor hDPP4. Experimental studies found no evidence of binding from HKU5 to hDPP4. Bt133 conserves all contact residues used by Ty-HKU4 to bind hDPP4 in 24 (marked with a pink asterisk). MERS uses four of the same contact residues^33^ than Ty-HKU4 (indicated by a blue triangle). HKU5, the only virus in the list unable to bind hDPP4, does not share any of the 8 contact residues from Ty-HKU4, and it shows several deletions at the RBM. **b.** Comparison of viruses related to LYRa3 within the Sarbecovirus subgenus.

LYRa3 and LYRa11 are phylogenetically related to SARS-CoV Tor2, and both share a high sequence identity (89.7%, 89.7% resp.) at the S gene. A multiple sequence alignment for related viruses within the Sarbecovirus subgenus shows that LYRa11 and LYRa3 are identical at the RBM except at residue H441. Twelve of the 17 contact residues that SARS-CoV Tor2 uses to bind to hACE2 are conserved (See Fig. 4b). In contrast, experimental studies from bat coronavirus ZC45^37^ did not find evidence of binding to hACE2^34,35^. When compared to SAR-CoV Tor2, ZC45 shows three deletions at the RBM and different amino acids at all locations of the contact residues of SAR-CoV Tor2, except for one at Y449. These results suggest that LYRa3 binds to hACE2.

### Molecular dynamics confirm binding of S to human receptors for coronaviruses Bt133 and LYRa3

Phylogenetic analysis, alignment results and the h-BiP score suggest that bat coronaviruses Bt133 and LYRa3 potentially bind to human receptors and are, therefore, candidates for host expansion to humans. We then used molecular dynamics simulations (MD) to validate binding in silico and to determine their contact residues. Three-dimensional structures of the receptor binding domain (RBD) bound to their corresponding human receptor for Lyra3 and Bt133 were obtained from crystal structure data and molecular modeling (See Methods).

The S protein of LYRa3 shares 99% identity with LYRa11 with one single point mutation at the RBD. Since LYRa11 is known to bind hACE2^34,35^, we expect LYRa3 to have similar binding properties. Results from three independent simulations showed that the average H-bond count between the RBD of LYRa3 and hACE2 was 5.1 (See Table 4). Similarly, the average number of H-bonds between the RBD of LYRa11 and hACE2 was 5.8 H-bonds. The difference in the number of H-bonds between LYRa3 and LYRa11 was not statistically significant (p-value: 0.2947), suggesting that they have comparable binding energies. The contact residues for LYRa3 were unknown. Therefore, we identified all of its contact residues during the course of the simulation (Supplementary Table 1). Our simulations revealed that contact residues G492, N477, T490, G486 and Y485 were present in at least 45% of the sampled conformations. A comprehensive list of all the bonds and their respective frequencies are available in Supplementary Table 1.

**Table 4:** Average H-bonds between RBD and human receptor.

| Virus | Sim1 | Sim2 | Sim3 | Average (n=3) | Std. error | p-value |
| --- | --- | --- | --- | --- | --- | --- |
| LYRa11 | 6.6 | 5.8 | 5.1 | 5.8 | 0.75 |  |
| LYRa3 | 4.3 | 5.5 | 5.5 | 5.1 | 0.73 | 0.2947 |
| Ty-HKU4 | 6.4 | 5.6 | 6.1 | 6.0 | 0.41 |  |
| Bt133 | 6.6 | 4.8 | 4.8 | 5.4 | 1.03 | 0.3780 |

Average number of H-bonds between the RBD and the human receptor by MD simulation (Sim1, Sim2, Sim3). The average was computed from all available sampled conformations after reaching equilibrium (4 ns). The average number of H-bonds of the three simulations and corresponding standard error is also shown. A t-test was performed to compare the means between LYRa11 and LYRa3 and between Ty-HKU4 and Bt133. The p-value corresponds to the two tailed test.

LYRa11 is phylogenetically related to SARS-CoV and there is experimental evidence of binding from both to human receptor hACE2. LYRa11 conserves 12 (out of 17) of the contact residues used by SARS-CoV^34, 35^ (marked with a pink asterisk). At the RBM, LYRa3 differs from LYRa11 only at H441, which is not a contact residue used by SARS-CoV. Experimental studies found no evidence of binding from ZC45 to hACE2. ZC45 conserves only 2 out of the 17 contact residues from SARS-CoV, and it shows several deletions at the RBM.

Next, we analyzed the interaction between the S protein of Bt133 and hDPP4. Despite containing 13 mutations in its RBD, alignment at the RBM showed that the contact residues of Ty-HKU4 are also present in Bt133. Results from three independent simulations showed that the average count of H-bonds was 5.4 for Bt133 and 6.0 for Ty-HKU4 (Table 4). The difference in average values was not statistically significant (p-value: 0.3780), suggesting that Bt133 and Ty-HKU4 have comparable binding energies when bound to hDPP4.

Our simulations confirmed all reported contact residues between Ty-HKU4 and hDPP4^24^ except for Q544. Our simulations also revealed contact residues N468, S465 and Y460, which have not been previously reported (Supplementary Table 2). Figure 5 shows contact residues for both Ty-HKU4 (a) and Bt133 (b). Only two contact residues were frequently observed in both viruses. A bond between E518 (magenta) in the RBD and Q344 in hDPP4, and between N514 (red) in the RBD and R317 in hDPP4, were present in at least 94% of the sampled conformations. Two additional contact residues were present in at least 50% of the sampled conformations from Ty-HKU4: K506 (purple) and K547 (orange). However, these two contact residues were present in less than 39% sampled conformations of Bt133. Instead, only one additional contact residue was present for Bt133 in more than 70% of the sampled conformations: Q515 (yellow).

**Fig. 5:**
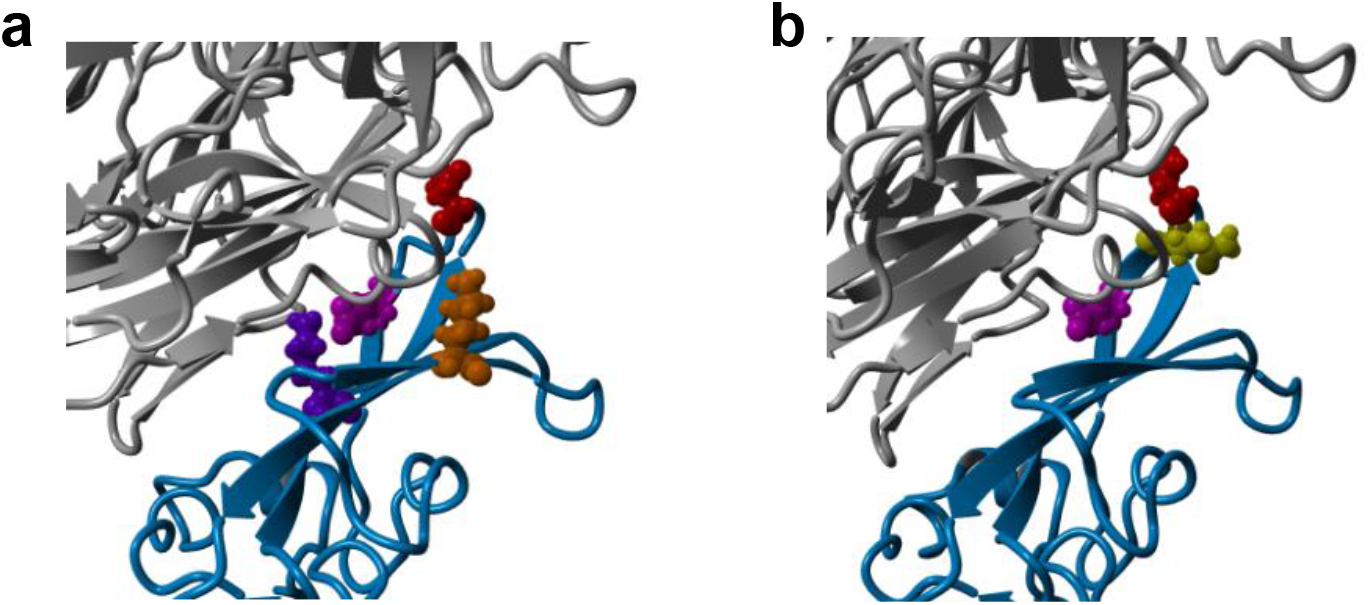
Most frequent contact residues for Ty-HKU4 and Bt133. The RBD is shown in light blue and the DPP4 human receptor in grey. Residues involved in frequent (average >=45% in Supplementary Tables 2 & 3) H-bonds are depicted in different colors. **a**. Frequent contact residues for Ty-HKU4 are E518 (magenta), N514 (red), K506 (purple) and K547 (orange). **b**. Frequent contact residues for Bt133 are E518 (magenta), N514 (red) and Q515 (yellow).

### The h-BiP score predicts binding of SARS-CoV2 to hACE2

We explored whether the proposed method can be used to detect potential human-infection of novel coronaviruses. To test this hypothesis, we repeated the study and computed a h-BiP score for SARS-CoV-2 after excluding all SARS-CoV-2 viruses from the training set. The new dataset consisted of 1,369 viruses with 540 labeled as positive for binding to a human receptor. The performance of the model in this subset reached 98% accuracy in the test set (sensitivity=99%, specificity=98%). The h-BiP score for the wild type of SARS-CoV-2 was 0.96; hence, the proposed method predicted binding of SARS-CoV-2 to a human receptor.

## Discussion

The COVID-19 pandemic has demonstrated the need to develop tools to predict spillover events. At the cellular and molecular level, three key steps, which are yet to be fully understood, are required for a spillover event: the evasion of the host’s immune system by the virus^4^, the infection of the cell by the virus^23,24^ and the replication of the virus in the new host cell^4,23^. In coronaviruses, the infection event is mediated by the binding of the viral spike (S) protein to the host receptor^23,24^. The vast abundance of recently collected S protein data opens the way for machine learning (ML) methods that will help accelerate the pace of discovery of virus candidates for spillover events.

Here, we propose a new machine learning approach to predict the binding of alpha and beta coronaviruses to a human receptor. We call this model the human-Binding Potential (h-BiP). As an alignment-free approach, h-BiP overcomes limitations associated with multiple sequence alignment (msa) methods, such as the lack of robustness against genomic rearrangements or low computational efficiency (O(nm) for msa)^19^. While most alignment-free approaches rely on k-mer counts, which disregards their relative position in the genome^14^, h-BiP uses a neural network to create k-mer embeddings through a context window. These embeddings encode information about their neighboring k-mers in the sequence, and the resultant classifier is highly accurate.

The h-BiP score is highly correlated with % identity against human coronaviruses. Yet, the classifier discriminates among viruses with similar % identity and identifies viruses with low % identity that may bind to human receptors. Such is the case of bat coronavirus LYRa3, which has a similar % identity to many bovine coronaviruses that do not bind human receptors; and the case of Bt133 with a low % identity (67.2%). However, they both have an h-BiP score greater than 0.5. The method suggests that these viruses with previously unknown human binding capabilities may bind to a human receptor. Phylogenetic analysis revealed that they are closely related to Ty-HKU4 and LYRa11, respectively, viruses known to bind human receptors. A pairwise alignment between the RBDs of Ty-HKU4 and Bt133 revealed that Bt133 conserves all 8 contact residues used by Ty-HKU4 to bind hDPP4 despite having 13 mutations in the RBD. On the other hand, with the exception of residue 441, the protein sequences of LYRa3 and LYRa11 are identical at the RBM. These findings suggest that Bt133 and LYRa11 bind to hDPP4 and hACE2, respectively.

We confirmed binding of Bt133 and LYRa3 in silico through a combination of structural modeling and molecular dynamics simulations. The average number of contact residues from the simulations of both viruses was not statistically different from that of Ty-HKU4 and LYRa11, respectively. These results suggest they have comparable binding energies. We identified all contact residues (Supplementary Table 2 and 3). Only contact residues E518 and N514 were frequently used (>94% of sampled conformations) by both viruses to bind to the human receptor. Contact residue E518 is also known to be relevant for binding of MERS^24,33^ to hDPP4. Our simulations also revealed three previously unreported contact residues from Ty-HKU4 (N468, S465 and Y460). Our study also identified the contact residues of LYRa3 (Supplementary Table 1).

As expected with any classifier, h-BiP produced a few false negatives in our dataset. Such is the case of bovine coronavirus isolate alpaca (h-BiP=0.4), which is the only known member from bovine coronaviruses experimentally confirmed for human binding in our dataset. However, no direct alpaca-human transmission has been reported. We hypothesize that a set of closely related sequences with mixed human receptor binding capabilities (positive and negative) produces scores near the classification threshold, thus reflecting the uncertainty of the prediction. Phylogenetic analysis under similar conditions would also lead to inconclusive findings.

We also tested the proposed method under the scenario for the emergence of a novel virus. In particular, we asked whether h-BiP would predict the binding of a novel coronavirus such as SARS-CoV-2. In the absence of all SARS-CoV-2 viruses, the h-BiP score for the wild type of SARS-CoV-2 was 0.96, demonstrating that h-BiP may be a valuable tool to detect the potential of a virus to cross species and originate an epidemic.

## Methods

### Datasets

On 11/05/2020, we downloaded 28,368 RNA spike protein sequences of all alpha and beta coronaviruses from the NCBI Virus^49^ database. Compared to the number of full nucleotide sequences, the number of annotated sequences for the Sarbecovirus genus (excluding SARS-CoV-2) was limited. On 07/26/2021, we downloaded all available nucleotide sequences and extracted the S protein (See section extracting the S protein from full sequences for details), expanding the data from 78 to 194 unique sequences from the Sarbecovirus genus (excluding SARS-CoV-2). To reduce the impact of an unbalanced dataset, we randomly removed 50% of SARS-CoV-2 (under-sampling), preserving reference sequences. The final alpha and beta coronavirus dataset consisted of 2,534 amino acid sequences. We curated the host field by combining information from isolation source, submission notes and the related publication. We also removed 7 viruses that were genetically modified (Accessions: FJ882951, FJ882957, HQ890538, FJ882942, HQ890534, MT782114, MT782115) as well as a draft virus from a pangolin (MT084071). Sequences in the data set were annotated as positive or negative for human binding. Human coronaviruses, and viruses from non-human animal hosts with published binding evidence to a human receptor, were labeled as positive for binding (n=1,705). Viruses from non-human hosts reported as unable to bind human receptors, and those for which the binding condition was unknown (n=782), were labeled as negative for binding (n=829). A comprehensive list of viruses with confirmed binding status is available in Table 1. The final dataset, according to binding status, is available in Table 5.

**Table 5:** Final dataset by binding condition to human receptor.

| Host | Negative/Unknown | Positive | Total |  |
| --- | --- | --- | --- | --- |
| Human | 0 | 1530 | 1530 | 60.4% |
| Non-human | 829 | 175 | 1004 | 39.7% |
| **Total** | **32.7%** | **67.3%** | **2534** |  |

### Extracting the S protein from full sequences

On 07/26/2021, we downloaded all available nucleotide (nt) sequences from the Sarbecovirus genus (excluding SARS-CoV-2) for a total of 643 sequences with at least 1603 nt of length. The S protein contains two subunits: the S1 subunit containing the receptor-binding-domain, and the S2 subunit, which mediates membrane fusion^50,51^. Prior to SARS-CoV-2 emergence, a highly conserved domain across all coronaviruses (SFIEDLLFNKVTLADAGF, NCBI accession cl40439) was found in the S2 subunit^51^. In order to extract the S protein from the full RNA sequence, we first translated it into the three possible reading frames. Next, we located the conserved domain by searching for the “SFIEDLLFN” motif (allowing for at most one substitution). The final putative S protein was the sequence enclosed by the start and stop codons, which contained the “SFIEDLLFN” motif. A total of 554 S proteins (194 unique) were found and included in the final dataset. We tested different motif lengths and number of substitutions to locate the conserved domain, but 9 amino acids, and 1 substitution, were found to be optimal.

### Training and test sets

The final dataset of 2,534 sequences was split into a training (85%) and test set (15%), stratified by the following groups: hCoV-OC43, hCoV-HKU1, MERS, SARS-CoV-1, SARS-CoV-2, hCoV-NL63, hCoV-229E, other MERS-related, other Sarbecovirus, other Betacoronavirus, porcine epidemic diarrhea virus, and other Alphacoronavirus.

### Sequence embedding

In order to generate n-dimensional vectors from the protein sequences, we first built “words” of 3 letters (trimers) with 3 contiguous amino acids. We considered all possible trimers by sliding a window one amino acid at a time as shown in Fig.1. To produce trimer embeddings, we used a skip-gram with negative sampling (vector-size=100, context-size=25, negative samples=1) as in 52. Finally, each sequence was represented as the sum of vectors from all of its trimers. That is, if a particular sequence (*seq_k_*) consisted of *n* trimers {*W_1_, W_2_, …, w_n_*}, and each trimer was represented by a 100-dimensional vector *W_i_*=[*x_i_1_, x_i_2_, …, x_i_100_*], then the final sequence embedding was calculated from equation (1).

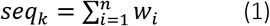

### Classification

In order to classify the sequences according to their potential to bind a human receptor, we labeled them as positive or negative for human binding as described in Datasets section. Normalized sequence embeddings of the training set were used to create a logistic regression model. The model used a ridge classifier with C=1.5 inverse regularization strength and the limited memory BFGS (L-BFGS) optimization algorithm. A virus was classified as positive for binding if the score was 0.5 or higher. The final classifier was invariant to the threshold value (see Supplementary Fig. 1).

### Phylogenetic tree

A phylogenetic tree for the S protein of alpha and beta coronaviruses was generated from a subset of 424 sequences (out of 2534) from the original dataset. This subset includes all reference viruses from both genera, several members from each of the 7 human coronaviruses and all viruses from non-human hosts with positive or negative published human binding annotation (See Table 1). A comprehensive list is available in Supplementary Table 4. A multiple sequence alignment (msa) of all sequences in the subset was generated with BBMap^53^. We used BEAST^54^ on the msa to reconstruct multiple phylogenetic trees with a log-normal distributed relaxed molecular clock and a non-parametric coalescent prior. The final tree shown in Supplementary Fig. 2 corresponds to the maximum-clade-credibility tree. We used iTOL^55^ for tree visualization and annotation.

### Candidate structures for molecular dynamics

The starting RBD structures for molecular dynamics (MD) simulations were obtained using PDB file 4QZV^24^ of Ty-HKU4-hDPP4 complex. At the RBD, Bt133 differs from Ty-HKU4 in 13 residues. These mutations were added to PDB 4QZV^24^ using YASARA^56^. The starting RBD structures for the RBD of LYRa3 and LYRa11 were generated using AlphaFold^57^ (implemented within the ColabFold suite^58^). Due to the absence of experimentally determined LYRa3-hACE2 and LYRa11-hACE2 complexes, we used the SARS-CoV-hACE2 complex (PDB 2AJF^43^) as a template in AlphaFold. The resulting structures were aligned to SARS-CoV using the MUSTANG^59^ method in YASARA^56^. These alignments were in strong agreement (<2.0A) and used as starting structures in our simulations.

### Molecular dynamics

To simulate protein-protein interactions, we used the molecular-modelling package YASARA^56^ to substitute individual residues and to search for minimum-energy conformations on the resulting modified candidate structures. For all structures, we carried out an energy minimization (EM) routine, which included steepest descent and simulated annealing minimization (until free energy stabilizes to within 50 J/mol) to remove clashes. All MD simulations were run using the AMBER14 force field^60^ for solute, GAFF2^61^ and AM1BCC^62^ for ligands and TIP3P^62^ for water. The cutoff was 8 Å for Van der Waals forces (AMBER’s default value^63^), and no cutoff was applied for electrostatic forces (using the Particle Mesh Ewald algorithm^64^). The equations of motion were integrated with a multiple timestep of 1.25 fs for bonded interactions and 2.5 fs for non-bonded interactions at T = 298 K and P = 1 atm (NPT ensemble) via algorithms described in 65. Prior to counting the RBD’s hydrogen bonds and calculating the free energy, we carried out several pre-processing steps on the structure, including an optimization of the hydrogen-bonding network^66^ to increase the solute stability and a pKa prediction to fine-tune the protonation states of protein residues at the chosen pH of 7.4^65^. Insertions and mutations were carried out using YASARA’s BuildLoop and SwapRes commands^65^, respectively.

Structure conformations from the simulations were collected every 100ps after 4ns of equilibration time as determined by the solute root mean square deviations (RMSDs) from the starting structure. For all bound structures, we ran the simulations for at least 10 ns post equilibrium and verified stability of time series for RBD-receptor hydrogen bond counts and root mean square deviation (RMSD) from these starting structures. Hydrogen bonds (H-bonds) were counted and tabulated using a distance and an angle approximation between donor and acceptor atoms as described in 66. It is important to note in this approach, salt bridges of proximate residues are effectively counted as H-bonds between basic side chain amide groups and acidic side chain carboxyl groups. Therefore, ionic interactions are also included in the H-bond count. In a previous publication, we show that the H-bond count, as defined here, correlates well with binding free energy estimates that were obtained using the molecular mechanics/generalized Born surface area method.

### Average number of H-bonds

In previous studies, we have shown that the average number of H-Bonds correlates well with binding energy^67^. Therefore, for each MD simulation, we recorded the number of H-bonds formed between protein-protein interactions (including ionic bonds) at each sampled conformation (snapshots). The average number of H-bonds was computed from all available sampled conformations after reaching equilibrium (4 ns). We performed three independent MD simulations from each complex and determined the grand average and standard error. Results are available in Table 4.

### H-bond frequencies (%)

At each sampled conformation (snapshot) from a MD simulation, we tracked every H-bond (including ionic bonds) and recorded the participant amino acids from the ligand and receptor. We computed the frequency for each pair dividing the number of sampled conformations where the pair was present by the total number of sampled conformations (double bonds are not considered in this count) and multiplied by 100. H-bond frequencies (%) from every simulation are available in the Supplementary Information.

## Data Availability

The datasets and corresponding GenBank accessions used to create h-BiP, together with the resultant scores, are available at https://github.com/Arsuaga-Vazquez-Lab/h-BiP.

## Code Availability

The h-BiP package and Python source code are available at https://github.com/Arsuaga-Vazquez-Lab/h-BiP.

## Acknowledgements

We thank Reuben Brasher for sharing best practices in repository management and helping with the testing platform. JA, MV, GG were partially supported by NSF grant DMS 2030491. GG was supported by a seed grant from CeDAR (Center for Data Science and Artificial Intelligence). GG was supported by a generous donation from Protein Architects to JA-MV group. We thank Michael Keith and Jacob Lusk for their valuable comments on the manuscript.

## Author contributions

G.G. J.A. and M.V. conceived the study design. G.G. downloaded and pre-processed the dataset. G.G. designed, coded, trained, tested and documented h-BiP. P.L. generated the phylogenetic tree. G.G. analyzed phylogenetic and h-BiP results. G.G. carried out sequence % identity and statistical analysis. M.J. and D.C. performed the structural modeling and the MD simulations. M.J., D.C. and G.G. analyzed the MD simulations. G.G., J.A. and M.V. wrote the manuscript with input from all other authors.

