## Supplemental Information for "Using machine learning to detect coronaviruses potentially infectious to humans"

Gonzalez-Isunza *et al.*

### Supplementary Information

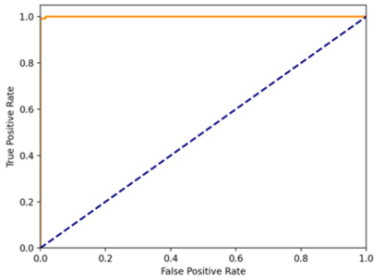


**Fig. 1: Receiver Operating Characteristic curve (ROC) from h-BiP scores.** The ROC curve on the alpha and beta coronaviruses test set at different thresholds of the h-BiP scores is shown in orange. The expected behavior of a random classifier is depicted with a blue dashed line. The area under the curve has a value of 0.999 showing that the performance of the model is invariant to the threshold's choice.

**a**

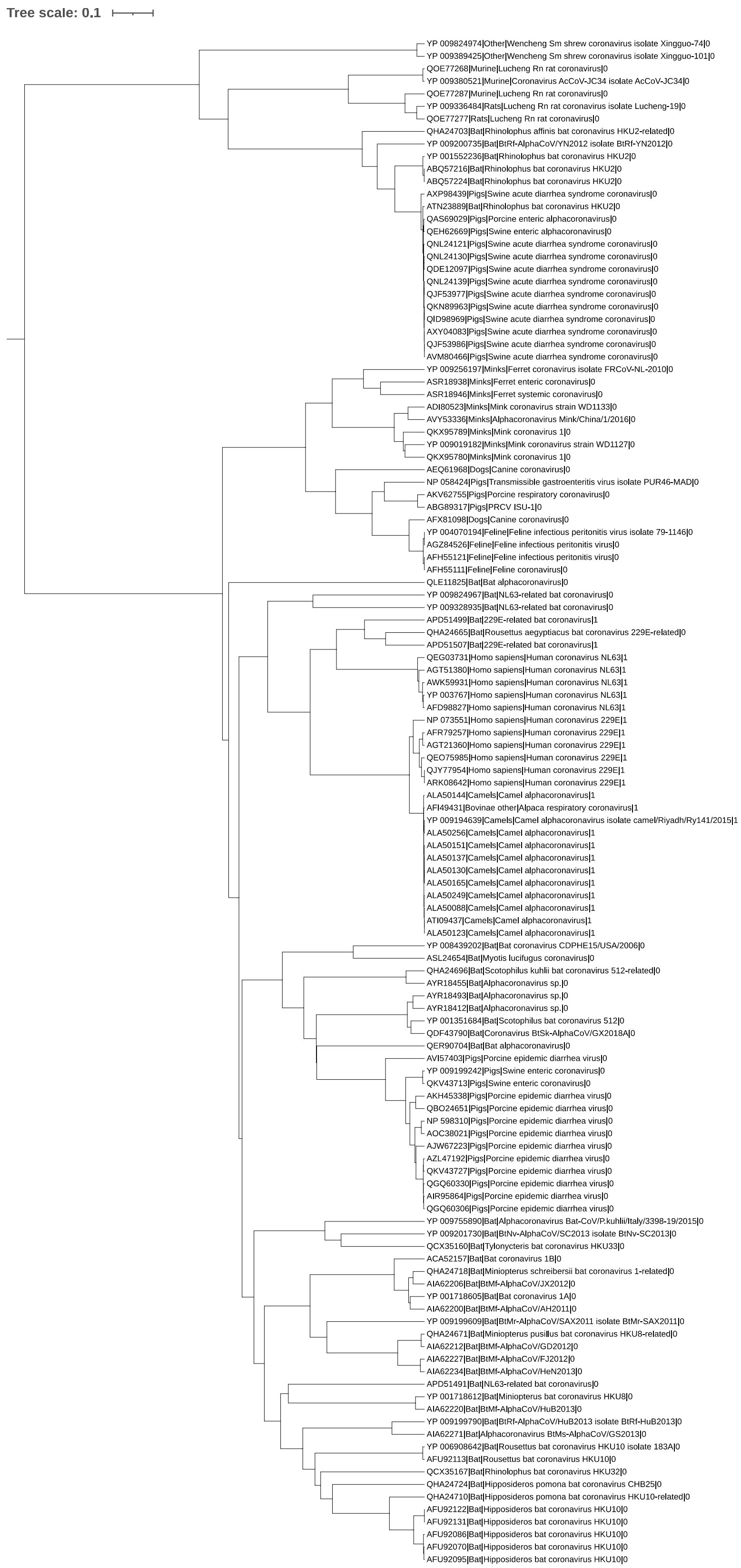

Tree scale: 0.1
YP 009824974|Other|Wencheng Sm shrew coronavirus isolate Xingguo-74|0
YP 009389425|Other|Wencheng Sm shrew coronavirus isolate Xingguo-101|0
QOE77268|Murine|Lucheng Rn rat coronavirus|0
YP 009380521|Murine|Coronavirus AcCoV-JC34 isolate AcCoV-JC34|0
QOE77287|Murine|Lucheng Rn rat coronavirus|0
YP 009336484|Rats|Lucheng Rn rat coronavirus isolate Lucheng-19|0
QOE77277|Rats|Lucheng Rn rat coronavirus|0
QHA24703|Bat|Rhinolophus affinis bat coronavirus HKU2-related|0
YP 009200735|Bat|BtRf-AlphaCoV/YN2012 isolate BtRf-YN2012|0
YP 001552236|Bat|Rhinolophus bat coronavirus HKU2|0
ABQ57216|Bat|Rhinolophus bat coronavirus HKU2|0
ABQ57224|Bat|Rhinolophus bat coronavirus HKU2|0
AXP98439|Pigs|Swine acute diarrhea syndrome coronavirus|0
ATN23889|Bat|Rhinolophus bat coronavirus HKU2|0
QAS69029|Pigs|Porcine enteric alphacoronavirus|0
QEH62669|Pigs|Swine enteric alphacoronavirus|0
QNL24121|Pigs|Swine acute diarrhea syndrome coronavirus|0
QNL24130|Pigs|Swine acute diarrhea syndrome coronavirus|0
QDE12097|Pigs|Swine acute diarrhea syndrome coronavirus|0
QNL24139|Pigs|Swine acute diarrhea syndrome coronavirus|0
QJF53977|Pigs|Swine acute diarrhea syndrome coronavirus|0
QKN89963|Pigs|Swine acute diarrhea syndrome coronavirus|0
QID98969|Pigs|Swine acute diarrhea syndrome coronavirus|0
AXY04083|Pigs|Swine acute diarrhea syndrome coronavirus|0
QJF53986|Pigs|Swine acute diarrhea syndrome coronavirus|0
AVM80466|Pigs|Swine acute diarrhea syndrome coronavirus|0
YP 009256197|Minks|Ferret coronavirus isolate FRCoV-NL-2010|0
ASR18938|Minks|Ferret enteric coronavirus|0
ASR18946|Minks|Ferret systemic coronavirus|0
ADI80523|Minks|Mink coronavirus strain WD1133|0
AVY53336|Minks|Alphacoronavirus Mink/China/1/2016|0
QKX95789|Minks|Mink coronavirus 1|0
YP 009019182|Minks|Mink coronavirus strain WD1127|0
QKX95780|Minks|Mink coronavirus 1|0
AEQ61968|Dogs|Canine coronavirus|0
NP 058424|Pigs|Transmissible gastroenteritis virus isolate PUR46-MAD|0
AKV62755|Pigs|Porcine respiratory coronavirus|0
ABG89317|Pigs|PRCV ISU-1|0
AFX81098|Dogs|Canine coronavirus|0
YP 004070194|Feline|Feline infectious peritonitis virus isolate 79-1146|0
AGZ84526|Feline|Feline infectious peritonitis virus|0
AFH55121|Feline|Feline infectious peritonitis virus|0
AFH55111|Feline|Feline coronavirus|0
QLE11825|Bat|Bat alphacoronavirus|0
YP 009824967|Bat|NL63-related bat coronavirus|0
YP 009328935|Bat|NL63-related bat coronavirus|0
APD51499|Bat|229E-related bat coronavirus|1
QHA24665|Bat|Rousettus aegyptiacus bat coronavirus 229E-related|0
APD51507|Bat|229E-related bat coronavirus|1
QEG03731|Homo sapiens|Human coronavirus NL63|1
AGT51380|Homo sapiens|Human coronavirus NL63|1
AWK59931|Homo sapiens|Human coronavirus NL63|1
YP 003767|Homo sapiens|Human coronavirus NL63|1
AFD98827|Homo sapiens|Human coronavirus NL63|1
NP 073551|Homo sapiens|Human coronavirus 229E|1
AFR79257|Homo sapiens|Human coronavirus 229E|1
AGT21360|Homo sapiens|Human coronavirus 229E|1
QEO75985|Homo sapiens|Human coronavirus 229E|1
QJY77954|Homo sapiens|Human coronavirus 229E|1
ARK08642|Homo sapiens|Human coronavirus 229E|1
ALA50144|Camels|Camel alphacoronavirus|1
AFI49431|Bovinae other|Alpaca respiratory coronavirus|1
YP 009194639|Camels|Camel alphacoronavirus isolate camel/Riyadh/Ry141/2015|1
ALA50256|Camels|Camel alphacoronavirus|1
ALA50151|Camels|Camel alphacoronavirus|1
ALA50137|Camels|Camel alphacoronavirus|1
ALA50130|Camels|Camel alphacoronavirus|1
ALA50165|Camels|Camel alphacoronavirus|1
ALA50249|Camels|Camel alphacoronavirus|1
ALA50088|Camels|Camel alphacoronavirus|1
ATI09437|Camels|Camel alphacoronavirus|1
ALA50123|Camels|Camel alphacoronavirus|1
YP 008439202|Bat|Bat coronavirus CDPHE15/USA/2006|0
ASL24654|Bat|Myotis lucifugus coronavirus|0
QHA24696|Bat|Scotophilus kuhlii bat coronavirus 512-related|0
AYR18455|Bat|Alphacoronavirus sp.|0
AYR18493|Bat|Alphacoronavirus sp.|0
AYR18412|Bat|Alphacoronavirus sp.|0
YP 001351684|Bat|Scotophilus bat coronavirus 512|0
QDF43790|Bat|Coronavirus BtSk-AlphaCoV/GX2018A|0
QER90704|Bat|Bat alphacoronavirus|0
AVI57403|Pigs|Porcine epidemic diarrhea virus|0
YP 009199242|Pigs|Swine enteric coronavirus|0
QKV43713|Pigs|Swine enteric coronavirus|0
AKH45338|Pigs|Porcine epidemic diarrhea virus|0
QBO24651|Pigs|Porcine epidemic diarrhea virus|0
NP 598310|Pigs|Porcine epidemic diarrhea virus|0
AOC38021|Pigs|Porcine epidemic diarrhea virus|0
AJW67223|Pigs|Porcine epidemic diarrhea virus|0
AZL47192|Pigs|Porcine epidemic diarrhea virus|0
QKV43727|Pigs|Porcine epidemic diarrhea virus|0
QGQ60330|Pigs|Porcine epidemic diarrhea virus|0
AIR95864|Pigs|Porcine epidemic diarrhea virus|0
QGQ60306|Pigs|Porcine epidemic diarrhea virus|0
YP 009755890|Bat|Alphacoronavirus Bat-CoV/P.kuhlii/Italy/3398-19/2015|0
YP 009201730|Bat|BtNv-AlphaCoV/SC2013 isolate BtNv-SC2013|0
QCX35160|Bat|Tylonycteris bat coronavirus HKU33|0
ACA52157|Bat|Bat coronavirus 1B|0
QHA24718|Bat|Miniopterus schreibersii bat coronavirus 1-related|0
AIA62206|Bat|BtMf-AlphaCoV/JX2012|0
YP 001718605|Bat|Bat coronavirus 1A|0
AIA62200|Bat|BtMf-AlphaCoV/AH2011|0
YP 009199609|Bat|BtMr-AlphaCoV/SAX2011 isolate BtMr-SAX2011|0
QHA24671|Bat|Miniopterus pusillus bat coronavirus HKU8-related|0
AIA62212|Bat|BtMf-AlphaCoV/GD2012|0
AIA62227|Bat|BtMf-AlphaCoV/FJ2012|0
AIA62234|Bat|BtMf-AlphaCoV/HeN2013|0
APD51491|Bat|NL63-related bat coronavirus|0
YP 001718612|Bat|Miniopterus bat coronavirus HKU8|0
AIA62220|Bat|BtMf-AlphaCoV/HuB2013|0
YP 009199790|Bat|BtRf-AlphaCoV/HuB2013 isolate BtRf-HuB2013|0
AIA62271|Bat|Alphacoronavirus BtMs-AlphaCoV/GS2013|0
YP 006908642|Bat|Rousettus bat coronavirus HKU10 isolate 183A|0
AFU92113|Bat|Rousettus bat coronavirus HKU10|0
QCX35167|Bat|Rhinolophus bat coronavirus HKU32|0
QHA24724|Bat|Hipposideros pomona bat coronavirus CHB25|0
QHA24710|Bat|Hipposideros pomona bat coronavirus HKU10-related|0
AFU92122|Bat|Hipposideros bat coronavirus HKU10|0
AFU92131|Bat|Hipposideros bat coronavirus HKU10|0
AFU92086|Bat|Hipposideros bat coronavirus HKU10|0
AFU92070|Bat|Hipposideros bat coronavirus HKU10|0
AFU92095|Bat|Hipposideros bat coronavirus HKU10|0

**b**

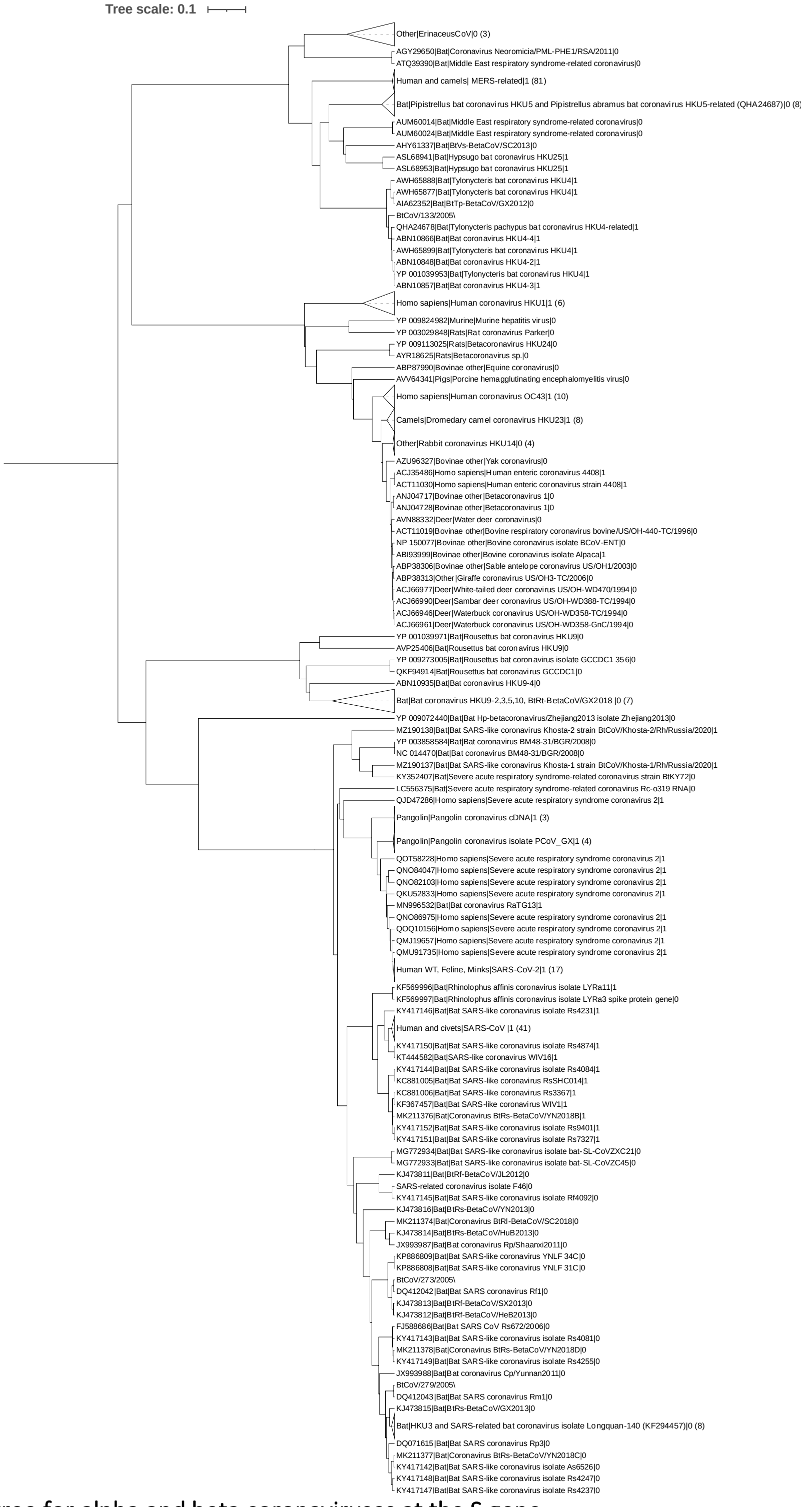


**Fig. 2: Phylogenetic tree for alpha and beta coronaviruses at the S gene**.
Amino acid sequences from the S gene of 424 alpha and beta coronaviruses (Supplementary Table 4) were aligned using BBMap. The phylogenetic tree was generated using maximum-clade-credibility tree with BEAST and visualized with iTOL. Each leaf shows the accession number from NCBI, the host, the GenBank Title and the label for the binding status separated by a pipe symbol (1 if there is experimental evidence of binding to human receptor, 0 otherwise). Collapsed clades show the total number of members in parenthesis **a.** Alphacoronavirus genus **b.** Betacoronavirus genus

Table 1: Hydrogen bond frequencies for all MD simulations of LYRa3-hACE2

| LYRa3 | hACE2 | Sim1 (n=160) | Sim2 (n=168) | Sim3 (n=122) | Average | Std. error |
|---|---|---|---|---|---|---|
| G492 | K353 | 92.5 | 94.0 | 98.3 | 94.9 | 2.5 |
| | G354 | 0.0 | 0.6 | 0.0 | 0.2 | 0.3 |
| N477 | Y83 | 89.3 | 98.2 | 90.1 | 92.5 | 4.0 |
| | Q24 | 33.3 | 40.5 | 57.0 | 43.6 | 9.9 |
| T490 | D355 | 63.5 | 41.7 | 62.0 | 55.7 | 10.0 |
| | Y41 | 10.7 | 25.6 | 1.7 | 12.6 | 9.9 |
| | N330 | 0.6 | 4.8 | 0.0 | 1.8 | 2.1 |
| G486 | K353 | 16.4 | 73.2 | 53.7 | 47.8 | 23.6 |
| Y485 | K353 | 35.2 | 63.7 | 43.0 | 47.3 | 12.0 |
| R430 | E329 | 34.6 | 10.1 | 43.8 | 29.5 | 14.2 |
| | Q325 | 1.9 | 0.6 | 0.0 | 0.8 | 0.8 |
| Y488 | Q42 | 22.6 | 19.6 | 16.5 | 19.6 | 2.5 |
| | D38 | 0.0 | 26.2 | 3.3 | 9.8 | 11.6 |
| Y495 | E37 | 10.1 | 17.9 | 0.8 | 9.6 | 7.0 |
| | R393 | 0.0 | 4.2 | 0.0 | 1.4 | 2.0 |
| Y479 | Q24 | 4.4 | 18.5 | 0.8 | 7.9 | 7.6 |
| | Y83 | 0.0 | 0.6 | 0.0 | 0.2 | 0.3 |
| N483 | K31 | 9.4 | 2.4 | 9.1 | 7.0 | 3.2 |
| | E35 | 0.6 | 0.6 | 4.1 | 1.8 | 1.7 |
| W480 | K31 | 10.1 | 0.0 | 9.1 | 6.4 | 4.5 |
| Y444 | H34 | 3.8 | 0.6 | 14.0 | 6.1 | 5.7 |
| N491 | K353 | 8.8 | 0.0 | 9.1 | 6.0 | 4.2 |
| | Y41 | 4.4 | 5.4 | 0.0 | 3.3 | 2.3 |
| S446 | K31 | 1.3 | 0.0 | 6.6 | 2.6 | 2.9 |
| D467 | Q24 | 2.5 | 0.0 | 0.0 | 0.8 | 1.2 |
| L482 | K31 | 0.6 | 0.0 | 1.7 | 0.8 | 0.7 |
| S437 | Q42 | 0.0 | 1.2 | 0.0 | 0.4 | 0.6 |
| Q496 | Q325 | 0.6 | 0.0 | 0.0 | 0.2 | 0.3 |
| | T324 | 0.0 | 0.6 | 0.0 | 0.2 | 0.3 |

Hydrogen bond frequencies (%) for three independent MD simulations of LYRa3 RBD bound to human receptor ACE2 (details in Methods). The number of sampled conformations is shown in parenthesis.

Table 2: Hydrogen bond frequencies for all MD simulations of Ty-HKU4-hDPP4

| Ty-HKU4 | hDPP4 | Sim1 (n=96) | Sim2 (n=171) | Sim3 (n=126) | Average | Std. error |
|---|---|---|---|---|---|---|
| E518 | Q344 | 93.8 | 96.5 | 95.2 | 95.2 | 1.1 |
| | A291 | 20.8 | 53.8 | 59.5 | 44.7 | 17.1 |
| N514 | R317 | 92.7 | 95.9 | 94.4 | 94.4 | 1.3 |
| | Y322 | 1.0 | 0.0 | 0.0 | 0.3 | 0.5 |
| K506 | A289 | 54.2 | 48.0 | 57.1 | 53.1 | 3.8 |
| | T288 | 34.4 | 31.6 | 32.5 | 32.8 | 1.2 |
| K547 | I295 | 50.0 | 49.7 | 50.8 | 50.2 | 0.5 |
| | L294 | 2.1 | 1.2 | 1.6 | 1.6 | 0.4 |
| E541 | K267 | 37.5 | 38.0 | 23.8 | 33.1 | 6.6 |
| N468 | R336 | 31.3 | 14.6 | 46.8 | 30.9 | 13.2 |
| | G335 | 0.0 | 0.0 | 0.8 | 0.3 | 0.4 |
| D542 | K267 | 12.5 | 29.2 | 28.6 | 23.4 | 7.7 |
| | Q286 | 6.3 | 0.6 | 0.0 | 2.3 | 2.8 |
| Q515 | R317 | 3.1 | 23.4 | 29.4 | 18.6 | 11.2 |
| | S292 | 44.8 | 0.0 | 0.0 | 14.9 | 21.1 |
| | Y322 | 0.0 | 2.3 | 0.8 | 1.0 | 1.0 |
| S465 | D331 | 0.0 | 2.3 | 51.6 | 18.0 | 23.8 |
| | S333 | 4.2 | 19.3 | 19.8 | 14.4 | 7.3 |
| | R336 | 0.0 | 12.3 | 0.0 | 4.1 | 5.8 |
| | S334 | 1.0 | 0.0 | 0.8 | 0.6 | 0.4 |
| D516 | Y322 | 38.5 | 0.6 | 1.6 | 13.6 | 17.7 |
| Y460 | G335 | 8.3 | 4.7 | 2.4 | 5.1 | 2.5 |
| | S333 | 0.0 | 0.0 | 3.2 | 1.1 | 1.5 |
| | S334 | 3.1 | 0.0 | 0.0 | 1.0 | 1.5 |
| | T283 | 0.0 | 1.2 | 0.0 | 0.4 | 0.6 |
| R462 | S333 | 0.0 | 11.7 | 0.0 | 3.9 | 5.5 |
| | E332 | 0.0 | 1.2 | 0.0 | 0.4 | 0.6 |
| C590 | N42 | 8.3 | 0.0 | 0.0 | 2.8 | 3.9 |
| S505 | R336 | 0.0 | 0.0 | 7.9 | 2.6 | 3.7 |
| D587 | D216 | 6.3 | 0.0 | 0.0 | 2.1 | 2.9 |
| | N40 | 4.2 | 0.0 | 0.0 | 1.4 | 2.0 |
| V580 | N109 | 4.2 | 0.0 | 0.0 | 1.4 | 2.0 |
| G543 | K267 | 0.0 | 0.6 | 1.6 | 0.7 | 0.7 |
| M592 | N44 | 2.1 | 0.0 | 0.0 | 0.7 | 1.0 |
| G585 | T213 | 2.1 | 0.0 | 0.0 | 0.7 | 1.0 |
| | S217 | 2.1 | 0.0 | 0.0 | 0.7 | 1.0 |
| G464 | S333 | 0.0 | 0.0 | 1.6 | 0.5 | 0.7 |
| A466 | S334 | 0.0 | 0.0 | 1.6 | 0.5 | 0.7 |
| G467 | S334 | 0.0 | 0.6 | 0.8 | 0.5 | 0.3 |
| S579 | N109 | 1.0 | 0.0 | 0.0 | 0.3 | 0.5 |
| T584 | S217 | 1.0 | 0.0 | 0.0 | 0.3 | 0.5 |
| S459 | S334 | 1.0 | 0.0 | 0.0 | 0.3 | 0.5 |
| S588 | D216 | 1.0 | 0.0 | 0.0 | 0.3 | 0.5 |
| Y472 | R336 | 0.0 | 0.0 | 0.8 | 0.3 | 0.4 |

Hydrogen bond frequencies (%) for three independent MD simulations of Ty-HKU4 RBD bound to human receptor DPP4 (details in Methods).The number of sampled conformations is shown in parenthesis.

Table 3: Hydrogen bond frequencies for all MD simulations of Bt133-hDPP4

| Bt133 | hDPP4 | Sim1 (n=135) | Sim2 (n=106) | Sim3 (n=113) | Average | Std. error |
|---|---|---|---|---|---|---|
| E518 | Q344 | 97.0 | 100.0 | 100.0 | 99.0 | 1.4 |
| | A291 | 51.1 | 16.0 | 14.2 | 27.1 | 17.0 |
| N514 | R317 | 96.3 | 95.3 | 95.6 | 95.7 | 0.4 |
| | Y322 | 0.7 | 0.0 | 0.0 | 0.2 | 0.3 |
| Q515 | S292 | 88.1 | 74.5 | 50.4 | 71.0 | 15.6 |
| | R317 | 5.9 | 0.0 | 0.9 | 2.3 | 2.6 |
| | Y322 | 0.0 | 0.9 | 1.8 | 0.9 | 0.7 |
| N468 | R336 | 64.4 | 49.1 | 16.8 | 43.4 | 19.8 |
| K506 | A289 | 57.8 | 0.9 | 57.5 | 38.7 | 26.7 |
| | T288 | 25.2 | 7.5 | 32.7 | 21.8 | 10.6 |
| K547 | I295 | 38.5 | 18.9 | 53.1 | 36.8 | 14.0 |
| | L294 | 5.2 | 16.0 | 0.0 | 7.1 | 6.7 |
| E541 | K267 | 21.5 | 3.8 | 4.4 | 9.9 | 8.2 |
| D542 | K267 | 19.3 | 0.9 | 8.0 | 9.4 | 7.5 |
| | Q286 | 2.2 | 9.4 | 8.8 | 6.8 | 3.3 |
| S465 | S334 | 7.4 | 7.5 | 0.0 | 5.0 | 3.5 |
| | S333 | 0.0 | 0.0 | 6.2 | 2.1 | 2.9 |
| Y460 | G335 | 3.7 | 0.9 | 1.8 | 2.1 | 1.2 |
| | S334 | 0.0 | 1.9 | 0.0 | 0.6 | 0.9 |
| S459 | S334 | 3.7 | 0.0 | 0.0 | 1.2 | 1.7 |
| C590 | N42 | 0.7 | 0.0 | 0.0 | 0.2 | 0.3 |
| D516 | Y322 | 0.7 | 0.0 | 1.8 | 0.8 | 0.7 |
| G585 | T213 | 0.0 | 0.9 | 0.0 | 0.3 | 0.4 |
| | S217 | 0.0 | 0.9 | 0.0 | 0.3 | 0.4 |
| P591 | N42 | 0.0 | 0.9 | 0.0 | 0.3 | 0.4 |
| R462 | S333 | 0.0 | 0.0 | 7.1 | 2.4 | 3.3 |
| G543 | K267 | 0.0 | 0.0 | 1.8 | 0.6 | 0.8 |

Hydrogen bond frequencies (%) for three independent MD simulations of Bt133 RBD bound to human receptor DPP4 (details in Methods).The number of sampled conformations is shown in parenthesis.

Table 4: Alpha and beta coronaviruses used to create the phylogenetic tree in Supplementary Fig. 2

| |
|---|
| Non-human Sarbecovirus (100) |
| MZ190138, MZ190137, MT799524, MT799523, MT799521, MT782114, MT072864, MT040336, MT040335, MT040334, MT040333, MN996532, MK211378, MK211377, MK211376, MK211374, MG772934, MG772933, LC556375, KY417152, KY417151, KY417150, KY417149, KY417148, KY417147, KY417146, KY417145, KY417144, KY417143, KY417142, KY352407, KU973692, KT444582, KP886809, KP886808, KJ473816, KJ473815, KJ473814, KJ473813, KJ473812, KJ473811, KF569997, KF569996, KF367457, KF294457, KC881006, KC881005, JX993988, JX993987, JX163927, JX163926, GQ153548, GQ153547, GQ153544, GQ153543, GQ153542, FJ959407, FJ588686, DQ648857, DQ648856, DQ514532, DQ514531, DQ514530, DQ514529, DQ514528, DQ412043, DQ412042, DQ084199, DQ071615, DQ022305, AY687372, AY687371, AY687370, AY687368, AY687365, AY687362, AY687361, AY687360, AY687359, AY687356, AY687355, AY687354, AY686863, AY627048, AY627047, AY627045, AY613952, AY613950, AY613948, AY572038, AY572037, AY572036, AY572034, AY545919, AY545915, AY515512, AY304489, AY304488, AY304486, NC_014470* |
| Human coronavirus SARS-CoV (4) |
| AY525636, AY278554, AY278489, AY274119* |
| SARS-CoV-2 from human host (12) |
| QOT58228, QOQ10156, QNO86975, QNO84047, QNO82103, QMU91735, QMJ19657, QMI94525, QKU52833, QKE51026, QJD47286, YP_009724390* |
| SARS-CoV-2 from mink host (8) |
| QNJ45226, QNJ45178, QNJ45142, QNJ45106, QJS39579, QJS39567, QJS39543, QJS39507 |
| SARS-CoV-2 from feline host (6) |
| QLC48479, QLC48467, QLC48455, QLC48443, QLC48419, QLC48407 |
| hCoVOC43 from human host (8) |
| QEG03794, AXX83375, AVR40342, ATP16767, AIV42005, AIV41987, AGT51561, YP_009555241* |
| hCoVOC43 from non-human host (2) |
| AWW13519, AWW13511 |
| Human coronavirus HKU1 (6) |
| AZS52618, ARB07438, AGW27872, ABD75601, ABD75513, YP_173238* |
| Human coronavirus NL63 (5) |
| QEG03731, AWK59931, AGT51380, AFD98827, YP_003767* |
| Human coronavirus 229E (20) |
| QJY77954, QEO75985, ATI09437, ARK08642, APD51507, APD51499, ALA50256, ALA50249, ALA50165, ALA50151, ALA50144, ALA50137, ALA50130, ALA50123, ALA50088, AGT21360, AFR79257, AFI49431, YP_009194639*, NP_073551* |
| MERS from human host (9) |
| QGW51920, QCQ29075, QBF80611, ASU45719, ANF29261, AKN24830, AKN24812, YP_009047204*, YP_007188579* |
| MERS from non-human host in the Middle East (66) |
| QOU08625, QOU08585, QOU08574, QOU08541, QOU08497, QCI31480, QCI31469, ASU91208, ASU91010, ASU90857, ASU90802, ASU90791, ASU90747, ASU90681, ASU90604, ASU90549, ASU90527, ASU90516, ASU90439, ASU90406, ASU90373, ASU90362, ASU90340, ASU90329, ASU90307, ASU90241, ASU90230, ASU90197, ASU90186, ASU90175, ASU90142, ASU90076, ASU90010, ASU89988, ASU89966, ASU89955, ASU89944, ASU45818, AQZ41296, ANI69922, ANI69900, ANI69889, ANI69878, ANI69835, ANI69824, ALL26409, ALL26396, ALA50067, ALA49957, ALA49803, ALA49671, ALA49660, ALA49649, ALA49473, ALA49462, ALA49451, ALA49396, ALA49374, ALA49363, ALA49352, ALA49341, AHY22565, AHY22555, AHY22545, AHX71946, AHX00711 |
| MERS from non-human host in Africa (11) |
| QGV13484, QBM11748, AXP07345, AVN89387, AVN89365, AVN89324, AUM60024, AUM60014, ATQ39390, AHY61337, AGY29650 |
| Porcine epidemic diarrhea virus (11) |
| QKV43727, QGQ60330, QGQ60306, QBO24651, AZL47192, AVI57403, AOC38021, AKH45338, AJW67223, AIR95864, NP_598310* |
| Betacoronavirus 1 (24) |
| AZU96327, AVV64341, AVN88332, ANJ04728, ANJ04717, ACT11019, ACJ66990, ACJ66977, ACJ66961, ACJ66946, ABP87990, ABP38313, ABP38306, QEY10657, QEY10649, QEY10641, QEY10633, QEY10625, AHN64783, AHN64774, ACT11030, ACJ35486, ABI93999, NP_150077* |
| Other Betacoronaviruses (13) |
| QDF43840, AYR18625, YP_009824982*, YP_009513010*, YP_009273005*, YP_009113025*, YP_009072440*, YP_005454245*, YP_003858584*, YP_003029848*, YP_001039971*, YP_001039962*, YP_001039953* |
| Other Alphacoronaviruses (39) |
| QLE11825, QKV43713, QER90704, QEH62669, QDF43790, AYR18493, AYR18455, AYR18412, AVY53336, AKV62755, AIA62271, AGZ84526, AFX81098, AFH55121, AFH55111, AEQ61968, ABG89317, YP_009824974*, YP_009824967*, YP_009755890*, YP_009389425*, YP_009380521*, YP_009336484*, YP_009328935*, YP_009256197*, YP_009201730*, YP_009200735*, YP_009199790*, YP_009199609*, YP_009199242*, YP_009019182*, YP_008439202*, YP_006908642*, YP_004070194*, YP_001718612*, YP_001718605*, YP_001552236*, YP_001351684*, NP_058424* |
| Other coronaviruses (80) |
| QOE77287, QOE77277, QOE77268, QNL24139, QNL24130, QNL24121, QKX95789, QKX95780, QKN89963, QKF94914, QJF53986, QJF53977, QID98969, QHA24724, QHA24718, QHA24710, QHA24703, QHA24696, QHA24687, QHA24678, QHA24671, QHA24665, QGA70702, QGA70692, QDE12097, QCX35167, QCX35160, QAS69029, AXY04083, AXP98439, AWH65932, AWH65921, AWH65910, AWH65899, AWH65888, AWH65877, AVP25406, AVM80466, ATN23889, ASR18946, ASR18938, ASL68953, ASL68941, ASL24654, APD51491, ALA50080, AIA62352, AIA62343, AIA62234, AIA62227, AIA62220, AIA62212, AIA62206, AIA62200, AFU92131, AFU92122, AFU92113, AFU92095, AFU92086, AFU92070, AFE48827, AFE48817, AFE48805, ADM33582, ADM33574, ADM33566, ADM33558, ADI80523, ACA52157, ABQ57224, ABQ57216, ABN10935, ABN10927, ABN10919, ABN10893, ABN10884, ABN10866, ABN10857, ABN10848, ABG47052 |

Accession numbers of the 424 alpha and beta coronaviruses used to create the phylogenetic tree in Supplementary Fig. 2. Reference sequences are marked with an asterisk. The total number of viruses for each cell is shown in parenthesis.
